# *HMGA1* zebrafish co-orthologue hmga1b can modulate p53-dependent cellular responses but is unable to control the alternative splicing of *psen1*

**DOI:** 10.1101/241927

**Authors:** Seyed Hani Moussavi Nik, Morgan Newman, Amanda Lumsden, Tanya Jayne, Michael Lardelli

## Abstract

The HIGH MOBILITY GROUP AT-HOOK 1 (HMGA1) family of chromatin-binding proteins plays important roles in cellular responses to low oxygen. HMGA1 proteins regulate gene activity both in the nucleus and within mitochondria. They are expressed mainly during embryogenesis and their upregulation in cancerous cells indicates poor prognosis. The human HMGA1a isoform is upregulated under hypoxia via oxidative stress-dependent signalling and can then bind nascent transcripts of the familial Alzheimer’s disease gene PSEN2 to regulate alternative splicing to produce the truncated PSEN2 protein isoform PS2V. Zebrafish where hmga1a expression is induced by hypoxia to control splicing of the *psen1* gene to produce the PS2V-equivalent isoform PS1IV. Zebrafish possess a second gene with apparent HMGA1 orthology, hmga1b. Here we investigate the predicted structure of Hmga1b protein and demonstrate it to be co-orthologous to human HMGA1 and most similar in structure to human isoform HMGA1c. We show that forced over-expression of either hmga1a or hmga1b mRNA can suppress the action of the cytotoxin hydroxyurea in stimulating cell death and transcription of the genes mdm2 and cdkn1a that, in humans, are controlled by p53. Our experimental data support an important role for HMGA1 proteins in modulation of p53-dependent responses and illuminate the evolutionary subfunctionalisation.

## INTRODUCTION

The protein HIGH MOBILITY GROUP AT-HOOK 1 (HMGA1) is notable for numerous reasons. It is the only protein to be exported from the nucleus to act in mitochondria ^1,2^ and its presence in mitochondria changes during the cell cycle ^2^. The protein is induced by the oxidative stress produced by hypoxia ^3,4^ and so is present during early embryogenesis in many vertebrate species. It can also be upregulated during cancer progression and is regarded as an indicator of poor prognosis ^5^.

Cancerous tumours present hypoxic environments in which HMGA1 may be activated. Therefore, there has been considerable interest in the possible interaction of this protein with the tumour suppressor p53, that is very commonly lost during oncogenesis ^6^. An inhibitory interaction between the HMGA1 and the pro-apoptotic functions of p53 has been reported although retraction of a paper central to this research area has led to uncertainty over these ideas ^7^.

The human *PRESENILIN* genes, *PSEN1* and *PSEN2* are loci for approximately 80% of the mutations causing early onset, dominantly-inherited, familial Alzheimer’s disease (on a population frequency basis ^8^). HMGA1 not only binds chromatin but has also been demonstrated to bind to *PSEN2* transcripts to control hypoxia-induced alternative splicing of this gene in humans. Binding of HMGA1 protein to *PSEN2* transcripts results in exclusion of exon 5 sequence leading to a frameshift and premature termination of the coding sequence ^9–11^. These alternative “PS2V” spliced transcripts are not completely removed by nonsense-mediated decay and so can be translated to produce a truncated form of PSEN2 protein that suppresses the unfolded protein response (UPR) to reduce the cell death that can be induced by hypoxia. PS2V is thus part of a negative feedback mechanism that limits UPR activation ^11,12^. We have previously demonstrated that this mechanism of HMGA1 action exists also in zebrafish but acts, instead, on the transcripts of the zebrafish’s *psen1-*orthologous gene, *psen1*, rather than its *PSEN2* orthologue. This is strong evidence that the hypoxia-induced, HMGA1-dependent alternative splicing of *PRESENILIN* transcripts predates the duplication and divergence of the common ancestor of the *PSEN1* and *PSEN2* genes early in vertebrate evolution ^3,11^. The subfunctionalisation of *PSEN1* and *PSEN2* that occurred after the duplication event presumably occurred after the divergence of the teleost and tetrapod evolutionary lineages over 400 million years ago ^13^ The HMGA1-controlled splicing mechanism was lost from the *psen2* gene lineage of teleosts and the *PSEN1* lineage of the tetrapods, including humans.

We have previously shown that, in zebrafish, the *hmga1a* gene is orthologous to human *HMGA1*, is upregulated in embryos and adult brains by hypoxia, and can drive formation of the zebrafish PS2V-equivalent isoform, PS1IV. However, zebrafish possess a second “co-orthologue” of *HMGA1* named *hmga1b* that has not been characterised. In this paper we analyse the putative structure of the *hmga1b* protein product, its phylogenetic relationship to other *HMGA1* genes, and its embryonic expression. *hmga1b* transcription is regulated by hypoxia but, unlike *hmga1a*, the protein product of *hmga1b* cannot regulate alternative splicing of *psen1* transcripts to produce isoform PS1IV. Nevertheless, and similar to *hmga1a*, it appears able to suppress the regulatory effects of p53 on target genes so as to inhibit cell death. Thus, HMGA1 proteins act at multiple levels to suppress the cell death caused by hypoxia and oxidative stress.

### Materials and Methods

#### Ethics

This work was conducted under the auspices of The Animal Ethics Committee of The University of Adelaide and in accordance with EC Directive 86/609/EEC for animal experiments and the Uniform Requirements for Manuscripts Submitted to Biomedical Journals.

#### Zebrafish husbandry and experimental procedures

*Danio rerio* were bred and maintained at 28°C on a 14 h light/10 h dark cycle. Embryos were collected from natural mating, grown in embryo medium (E3) and staged ^14^. Treatment of embryos at 48 hours post fertilisation (hpf) and adult zebrafish with low oxygen and mimicry of hypoxia using sodium azide (NaN_3_) has been described previously ^3^. For treatment with hydroxyurea (HU), embryos were divided into two groups after 24 hpf and hydroxyurea was added to one of the two groups to a concentration of 250 mM. Incubation of both groups then continued until the developmental stage equivalent to that of untreated embryos at 48 hpf (at 28.5°C). Embryos were then processed for further analysis.

#### Phylogenetic analyses

Multiple alignment of HMGA-related proteins in human and zebrafish was done using ClustalW with the gap opening penalty of 10.0 and gap extension penalty of 3.0 ^15^. Phylogenetic analysis was conducted using MrBayes ^16^, the program was run under the JC69 (Juke-Cantor) model, with rate variation gamma and unconstrained branch lengths of 10.

#### Injection of mRNAs

Embryos were collected from natural mating, grown in embryo medium (E3) and staged. *In vitro*-synthesised mRNAs were injected into one-cell stage embryos as previously described^17^.

#### Quantitative real-time PCR assay

Total RNA was extracted from adult brain or embryos using the QIAGEN RNeasy mini kit (QIAGEN, GmbH, Hilden, Germany). cDNA synthesis, quantitative real-time PCR (qPCR) assay, and data analysis has been described previously ^11^. Gene specific primers used for qPCR are given in Supplementary Data File 2.

#### Western immunoblot analyses

At 48 hpf embryos were dechorionated, deyolked and placed in lithium dodecyl sulfate (LDS) sample buffer (NuPAGE^®^ LDS sample buffer 4X, NuPAGE^®^ Reducing reagent 10X), heated immediately at 80°C for 25 min and then stored at −80°C prior to separation on a 4–12% NuPAGE^®^ Gel. Western blotting and transfer of protein onto nitrocellulose membranes was performed as previously described ^17^. For incubation with anti-Cleaved Caspase 3 (Asp175) antibody (Cell Signaling Technology Inc., Danvers, MA), primary antibody was diluted 1 in 1000 in TBST containing 2% w/v skim milk. The membranes were then washed in TBST and incubated with a goat anti-rabbit IgG secondary antibody (Cell Signaling Technology Inc., Danvers, MA.) diluted 1 in 3000 in TBST containing 2% w/v skim milk. After incubation with the secondary antibody the membranes were washed and visualised as previously described ^17^. Incubation and detection of anti-β-tubulin antibodies (Antibody E7, Developmental Studies Hybridoma Bank, The University of Iowa, IA, USA) was done as previously described ^17^.

#### Assessment of relative cell death

To visualise and quantify the relative cell death produced by hydroxyurea treatment in 48 hpf embryos, acridine orange (AO) assays were performed as previously described ^3^ except that quantification was performed on the tail region of 6 embryos per treatment (to avoid a signal from the developing lens).

#### Promoter analysis

Genomatix Gene2Promoter software (Genomatix AG, Munich, Germany) was used to extract the promoter sequences of human *HMGA1*, mouse *Hmga1*, bovine *HMGA1*, chicken *HMGA1*, and zebrafish *hmga1a* and *hmga1b*. Sequences consisted of 1000 bp upstream of the first transcription start site (TSS) to 100 bp downstream of the last TSS in each analysed promoter region. Sequences were examined using the *CommonTFs* function of MatInspector software (Genomatix AG ^18^) to identify transcription factor matrix families (Vertebrate and General Core Promoter Elements) common to *HMGA1* gene orthologues from human, mouse, cow and chicken, and at least one zebrafish paralog. Default stringency conditions were used (Minimum Matrix Core similarity, 0.075; Minimum matrix similarity, ‘Optimized’). Matches with a significant frequency of occurrence (*p*-value less than 0.05, based on expected frequency in promoter sequences) were extracted and presented, with position within the extracted sequence (in the DNA strand phase of the gene) given.

### Results

#### Confirmation of zebrafish *hmga1b* as a co-orthologue of human *HMGA1*

We previously identified a zebrafish orthologue of the human *HMGA1* gene, at that time named *hmga1* ^3^. However, subsequent examination of zebrafish genome sequence led to putative identification of a second *HMGA1* “co-orthologous” gene, a common situation in zebrafish genetics ^19^. This second gene is now named *hmga1b* while the previously characterised gene is now *hmga1a*. The *hmga1b* gene is located on zebrafish chromosome 6 (RefSeq NM_ 001077276.1).

The human *HMGA1* gene expresses three protein isoforms (derived from alternative transcript splicing), HMGA1a, HMGA1b, and HMGA1c. These differ in the spacing and number of their DNA-binding AT-hook domains (see Figure 1A, B). Overall, the amino acid (aa) residue identity between the human HMGA1 and the putative protein sequence of zebrafish Hmga1b is 59 %. The two most N-terminal AT-hook domains in Hmga1b share identical aa sequence with human HMGA1 and zebrafish Hmga1a. However, while the most C-terminal AT-hook domain is also identical between human HMGA1 and zebrafish Hmga1a, this domain shows divergence in zebrafish Hmga1b. Hmga1b also apparently lacks the acidic tail domain that is conserved in zebrafish Hmga1a and human HMGA1a and HMGA1b (although not in isoform HMGA1c, see ^20^, Figure 1A, B).

**Figure 1.**
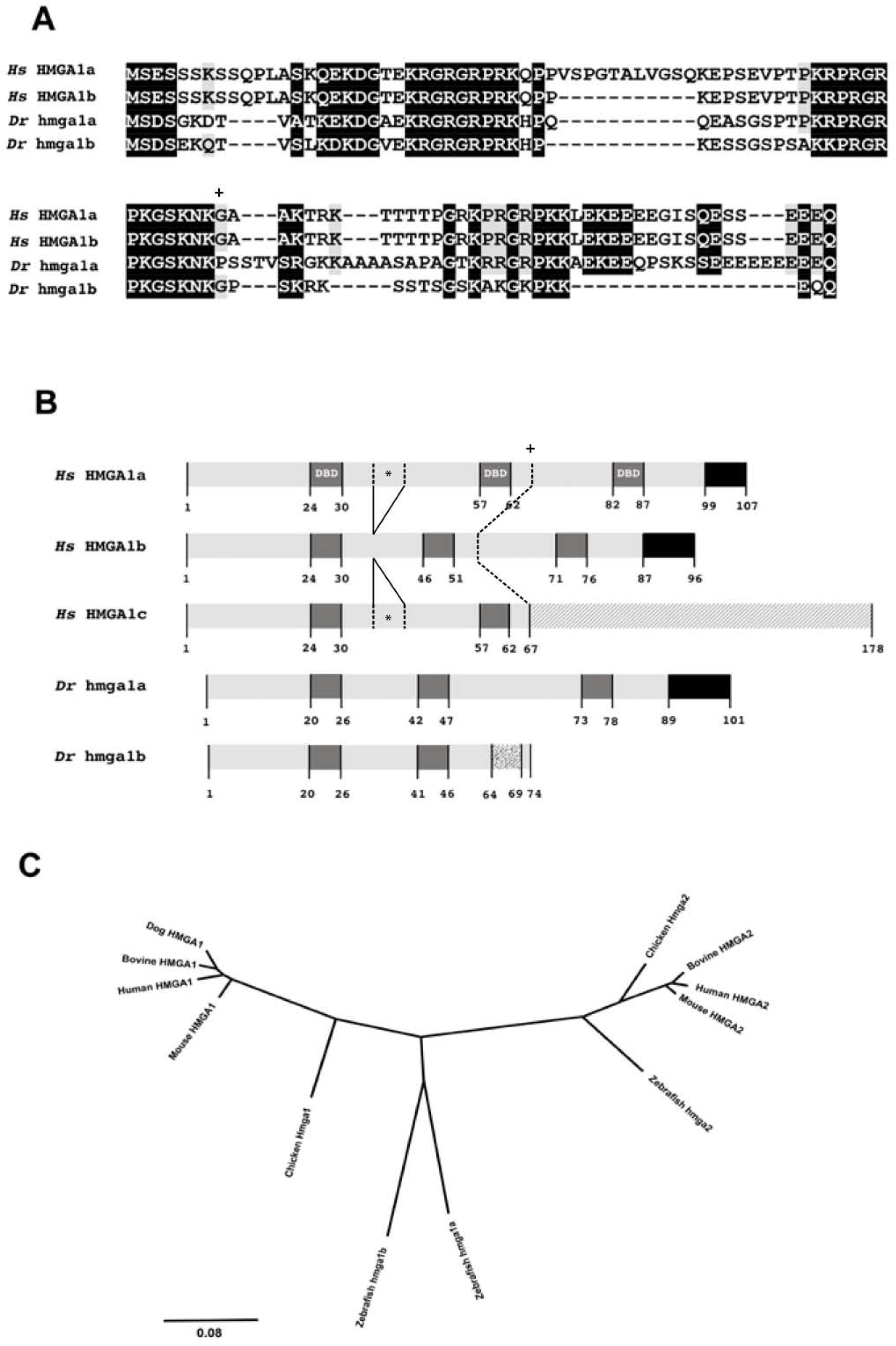
A) Amino acid residue sequence alignment of the zebrafish Hmga1b putative protein sequence and human HMGA1 performed using ClustalW. B) Schematic illustration comparing the structural differences between different HMGA1 proteins in Human (Hs) and zebrafish *(Dr)*. The dark grey shadinged areas indicate the AT-hook DNA binding domains (DBD) characterised by the sequence **K/R(X)RGRP**. The black shaded regions indicate the acidic tails of the proteins. A less conserved AT-hook-like domain near the C-terminal end of Hmga1b is indicated in stippled light grey. Additional protein sequence from an additional exon included in HMGA1a and HMGA1c is indicated by *. Sequence divergence between HMGA1c and the other HMGA1 isoforms due to alternative splicing is indicated by dotted lines and +. Numbers indicated amino acid residues. C) Phylogenetic tree of the high mobility group A protein family generated using MrBayes. Sequences used in the phylogenetic analysis are shown in Table S1 in Supplementary Data File 1.

To confirm the orthology relationships between human *HMGA1* and zebrafish *hmga1a* and *hmga1b*, we conducted a phylogenetic analysis comparing the zebrafish genes with DNA sequences of the *HMGA1* orthologues of human, mouse, bovine and chicken, as well as the most closely related *HMGA* gene family, the *HMGA2* genes, of these species (Figure 1C). (See Table S1 in Supplementary Data File 1 for sequence accession numbers and description of the exact sequences from the multiple cDNA sequence alignment used in the analysis). The close association between zebrafish *hmga1a* and *hmga1b* strongly supports that these two genes are co-orthologues of human *HMGA1*.

#### Expression of zebrafish *hmga1b*

The temporal expression of zebrafish *hmga1b* transcripts was observed by qRT-PCR analysis (relative to expression of housekeeping gene *eef1a111*) using three independent biological replicates for each developmental time point (the qPCR raw data and statistical analysis is available in Supplementary Data File 2). Particular adult tissues were also examined. Zebrafish *hmga1b* is present in embryos at two hours post fertilisation (hpf), before the mid-blastoderm transition when the zygotic genome becomes transcriptionally active ^21^. This indicates a maternal contribution of *hmga1b* mRNA to the embryo. We observed a peak of *hmga1b* mRNA levels at 48 hpf when embryogenesis ends with hatching, followed by a decline over subsequent days (Figure 2). Expression of *hmga1b* transcripts in adult brain, liver and kidney appeared lower than in whole animals up to 120 hpf (Figure 2).

**Figure 2.**
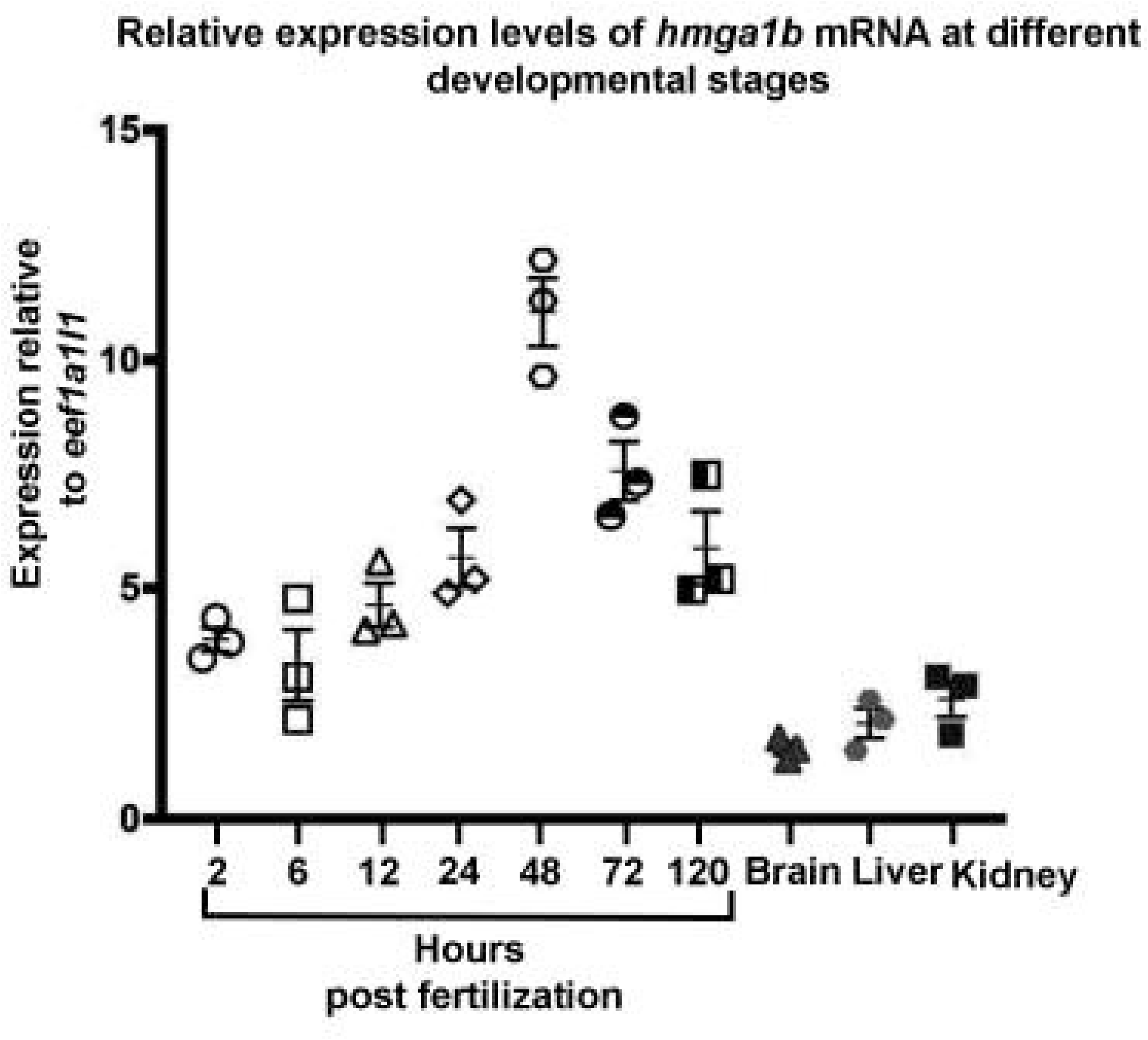
Temporal and tissue variation in expression of zebrafish *hmga1b* transcripts. Analysis of *hmga1b* expression by qPCR at various developmental time-points and in various adult tissues relative to housekeeping gene *eef1a111*. Means and standard error of the means are shown for three biological replicates.

#### Common hypoxia-related regulatory elements in promoters of HMGA1 homologs

The genes *hmga1a* (on chromosome 23) and *hmga1b* (on chromosome 6) probably arose in the teleost lineage-specific whole genome duplication that occurred over 300 million years ago ^22^. This is supported by the synteny of each gene with other genes that show similarity between the two chromosome regions. For example, the genes *nudt3a* and *pacsin1a* lie close downstream of *hmga1a* while genes *nudt3b* and *pacsin1b* are close downstream of *hmga1b* (Ensembl release 90, zebrafish GRCz10^23^).

After a gene duplication event, subfunctionalisation may occur where one of the duplicates uniquely retains particular activities of the ancestral gene while these are lost from the other duplicate copy (and vice-versa ^24^) The duplicate genes may also show some redundant function. Human *HMGA1* is both an important regulator of other genes and shows strong regulatory responses itself. For example, *HMGA1* is commonly upregulated in oncogenesis and by hypoxia ^25^. To identify transcription factor binding sites (TFBSs) that may be functionally relevant in the regulation of expression of HMGA1 homologs, we performed a comparative promoter analysis to find TFBS motifs common to a range of mammalian and avian *HMGA1* orthologues (human, mouse, bovine, chicken) that were also present in one or both of the zebrafish paralogs *hmga1a* and *hmga1b*. Promoter regions spanning 1000 nucleotides (nt) upstream of the first transcription start site (TSS) to 100 nt downstream of the last TSS in the analysed region were compared using the *CommonTFs* function of the program Genomatix MatInspector ^18^ to identify common transcription regulatory motifs. 28 matches were identified with a statistically significant frequency within the sequences (*p*-value <0.05; Table 2). All sequence positions and match scores of the regulatory motifs are presented in Supplementary Data File 2. Amongst the transcription factor binding motifs common to all the analysed homologs sites were found for factors involved in the hypoxic response ^26^ such as hypoxia inducible factor HIF family members (V$HIFF), and nuclear factor kappa B factors (V$NFKB) (Figure 3).

**Table 1:**
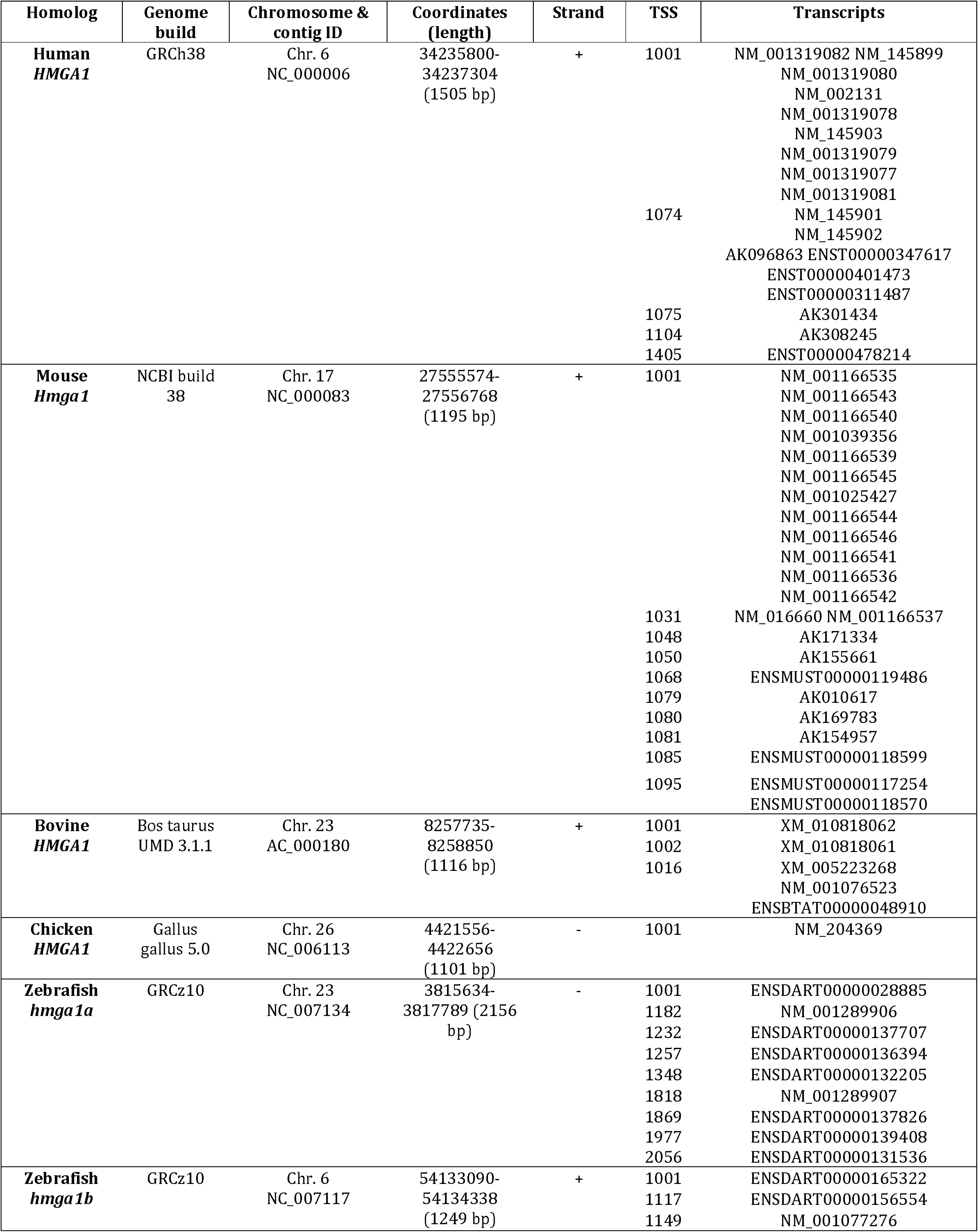
Genomic co-ordinates and associated transcripts of the promoter sequences of *HMGA1* homologs used for comparative analysis of transcription factor binding sites.

**Table 2:**
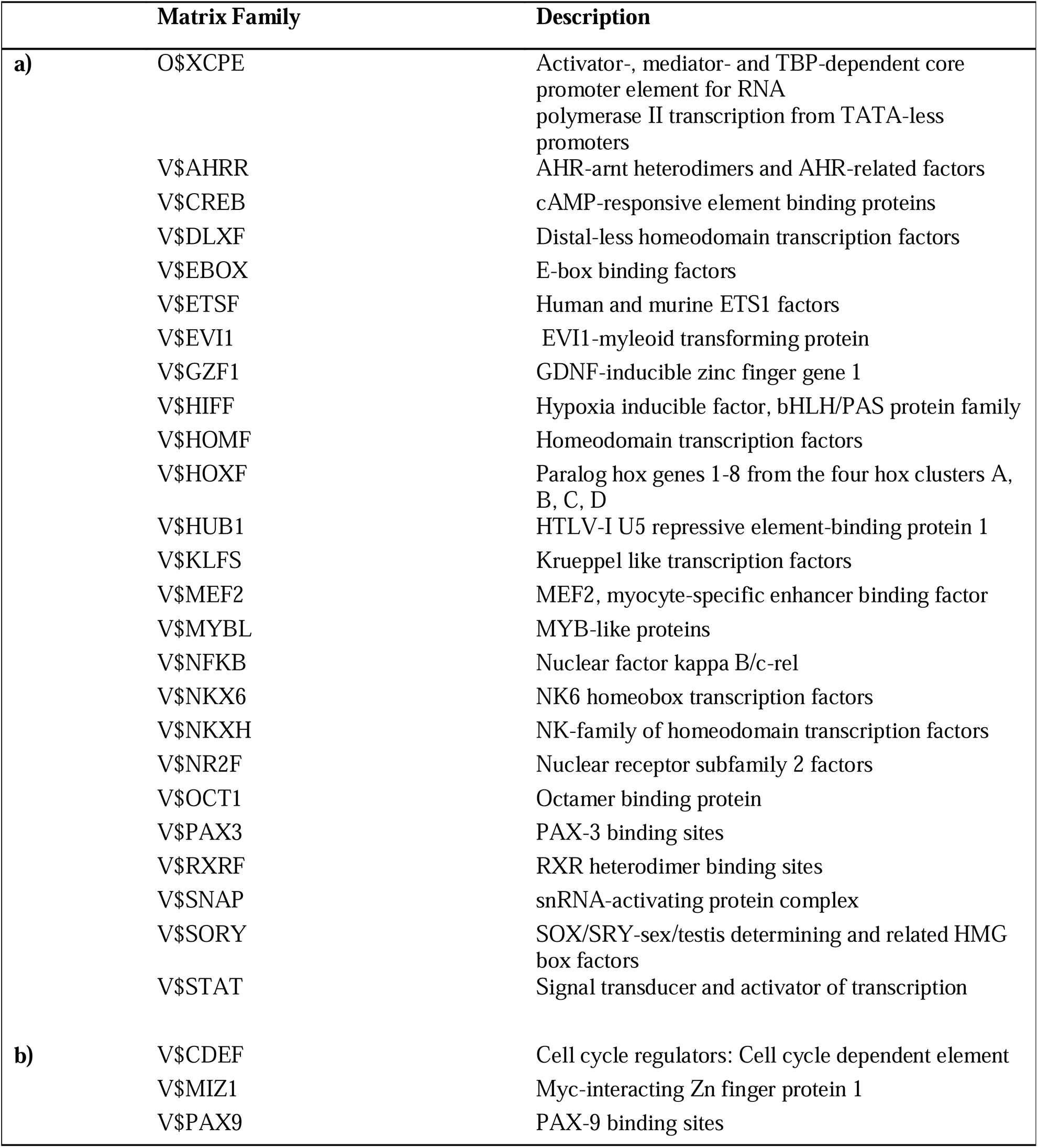
Transcription Factor Matrices present in the *HMGA1* promoter from human, mouse, bovine and chicken, that are also present in **a)** zebrafish *hmga1a* and *hmga1b* promoter sequences, or **b)** zebrafish *hmga1a* but not *hmga1b*. In this analysis, no sites common to human, mouse, bovine and chicken *HMGA1 were* detected in *hmga1b* that were not also found in *hmga1a*..

**Figure 3.**
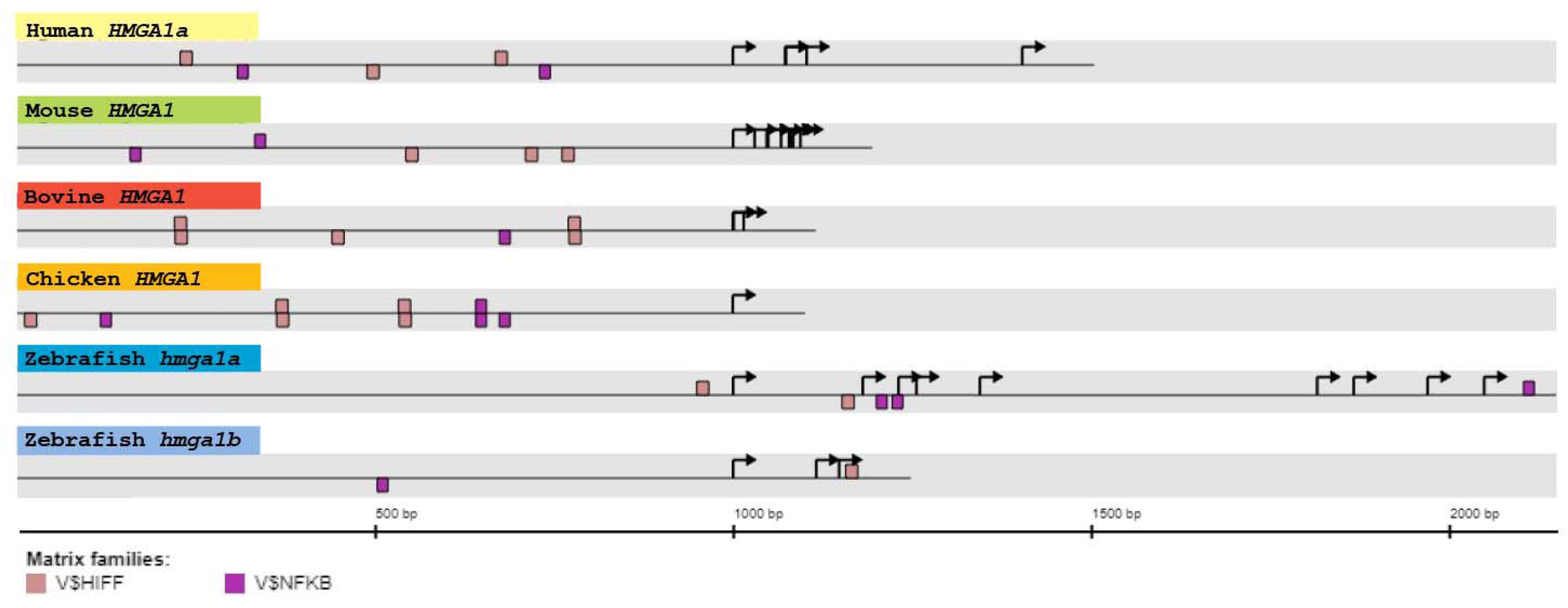
Location of transcription factor binding sites related to hypoxia response, within a selection of vertebrate *HMGA1* orthologue promoters. Genomatix schematic output modified for clarity. Genomic sequence coordinates and associated transcripts are presented in Table 1.

#### Testing regulation by hypoxia of zebrafish *hmga1b* transcript levels

In humans, hypoxia increases expression of *HMGA1* transcripts in neural tissue through induction of oxidative stress ^9,27^ and this explains the induction by hypoxia of the PS2V alterative transcript isoform from the *PSEN2* gene ^9^. We previously demonstrated that this mechanism is conserved in zebrafish with increased expression of *hmga1a* transcripts causing induction of the PS2V-cognate isoform PS1IV from the zebrafish *psen1* gene ^11^. In humans, the HMGA1a isoform of HMGA1 shows a greater ability to induce PS2V formation than HMGA1b ^10^ while the activity of isoform HMGA1c has not been tested in this regard. Structurally, the putative zebrafish Hmga1b protein resembles somewhat human HMGA1c in that Hmga1b possesses only two highly conserved AT-hook domains and lacks the long acidic C-terminal domain seen in HMGA1a, HMGA1b and zebrafish Hmga1a. Therefore, we examined experimentally whether hypoxia might also regulate expression of *hmga1b* and, if so, whether *hmga1b* expression could (like *hmga1a*) influence alternative splicing of *psen1* to form PS1IV.

Hypoxia (or chemical mimicry of hypoxia using NaN3) increases the expression of zebrafish *hmga1a* transcripts both in whole embryos and adult brain tissue ^3^. Therefore, we compared the relative levels of *hmga1b* mRNA in 48 hpf embryos under normoxia and chemical mimicry of hypoxia using qRT-PCR. Similarly, we examined the relative levels of *hmga1b* transcripts in the brains of adult zebrafish that had been exposed to normoxia or hypoxia. Under chemical mimicry of hypoxia or real hypoxia, relative induction of *hmga1b* (compared to transcript levels of the housekeeping gene *eef1a111*) is observed in 48 hpf embryos (Figure 4A) and adult brains (Figure 4B) respectively, to a degree similar to that of *hmga1a*.

**Figure 4.**
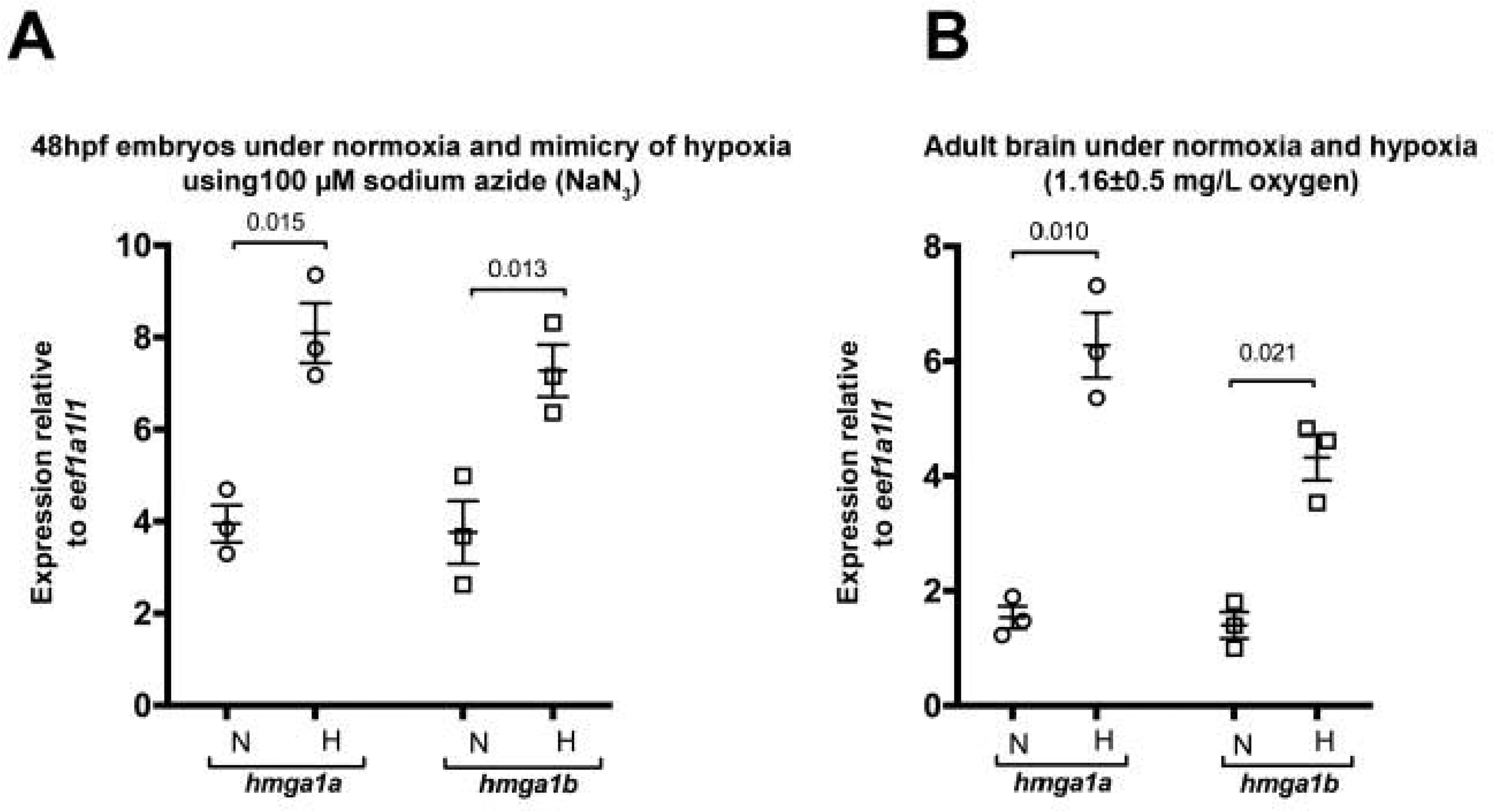
Response of *hmga1b* expression to **(A)** mimicry of hypoxia in embryos and **(B)** hypoxia in adult brain compared to *hmga1a* expression responses. qPCR analysis of *hmga1b* and *hmga1a* expression relative to eukaryotic translation factor *eef1a111*. Only relative responses to hypoxia, not absolute expression levels, can be compared between the two genes. Means and standard error of the mean are shown for three biological replicates (*n*=3). *P-*values for two-tailed T-tests (unequal variances) for bracketed comparisons are shown. N, Normoxia. H, Hypoxia.

#### Foreced over-expression of zebrafish *hmga1b* has no effect on zebrafish *psen1* alternative splicing

Forced embryonic expression of zebrafish *hmga1a* after injection of its mRNA into zebrafish zygotes causes increased alternative splicing of endogenous *psen1* transcripts to form PS1IV ^11^. Therefore, we compared this ability of *hmga1a* mRNA with that of *hmga1b* mRNA to test whether the latter can also control *psen1* alternative splicing. We used qPCR to compare levels of the PS1IV form of *psen1* mRNA in the presence of an injected, non-translated mRNA (as a negative control) or *hmga1a* mRNA or *hmga1b* mRNA. Only injection of *hmga1a* mRNA increased the level of PS1IV (Figure 5). Thus, only *hmga1a* appears able to regulate PS1IV production.

**Figure 5.**
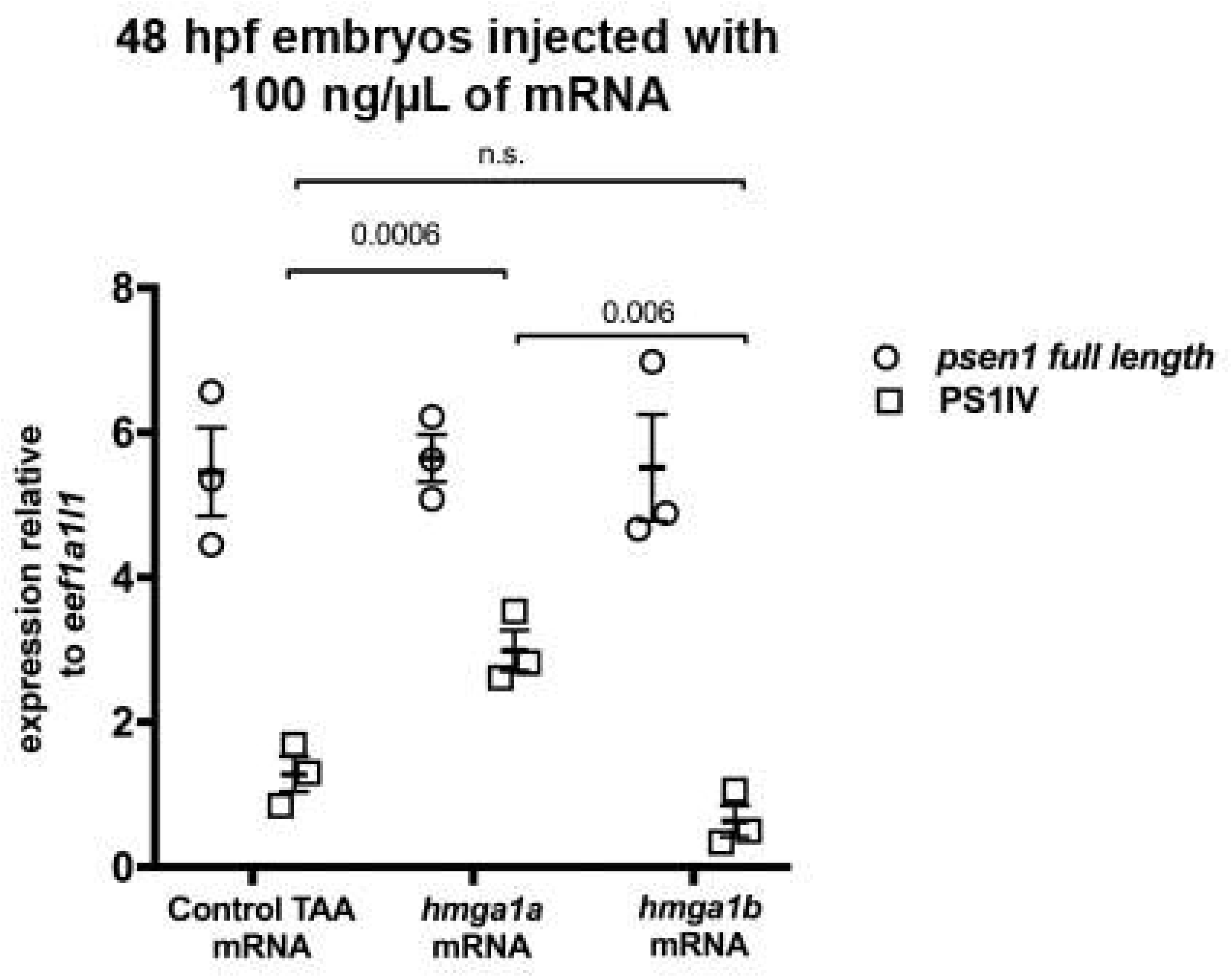
Injection of embryos with *hmga1a* mRNA, but not *hmga1b* mRNA, induces the formation of the *PSEN1* transcript isoform PS1IV under normoxia. qPCRs detecting, specifically, full-length *PSEN1* RNA or the alternatively-spliced PS1IV transcript isoform indicate induction of PS1IV by forced expression under normoxia of zebrafish Hmga1a, but not Hmga1b. Note that only comparisons of responses to forced expression of the same transcripts (i.e. full-length or PS1IV) are valid. Expression data is not comparable between full-length *PSEN1* and PS1IV. Expression of *PSEN1* and PS1IV is shown relative to eukaryote translation factor gene *eef1a111*. Means and standard error of the means are shown for three biological replicates in each case (*n*=3). *P*-values for two-tailed T-tests (unequal variances) for bracketed comparisons are shown. n.s., not statistically significant. Control TAA mRNA is an untranslatable control mRNA. No significant difference in levels of full-length *PSEN1* mRNA was observed for any treatment.

#### Zebrafish *hmga1b* decreases P53-dependent apoptosis in embryos

Upregulation of HMGA1 expression is a marker of poor prognosis in cancer ^28^. Research into the promotion of cancer progression by HMGA1 has identified that this protein may inhibit, by various means, the function of the tumour suppressor p53 (e.g. ^29^) although the research on this topic has been afflicted by controversy (e.g. see ^7^). Therefore, we examined whether the *hmga1a* and *hmga1b* genes of zebrafish might show activity supporting an interaction with p53-mediated cell death. The external development of zebrafish embryos makes them highly amenable to studying the effects of drug exposure and the cytotoxin hydroxyurea inhibits DNA synthesis and can induce apoptosis via a p53-dependent pathway ^30,31^. Therefore, we exposed embryos to 250 mM hydroxyurea to induce cell death and observed whether forced expression of *hmga1a* or *hmga1b* mRNA could reduce this.

Zygotes were injected with either an untranslatable mRNA (as a negative control) or with mRNA encoding proteins Hmga1a or Hmga1b before exposure to hydroxyurea at 24 hpf. At 48 hpf, embryos were then stained with acridine orange to visualise and quantify cell death (Figure 6A). Protein lysates from similarly-treated embryos were examined by western immunoblotting to quantify the levels of cleaved Caspase3 protein that is a marker of apoptosis (Figure 6B). We observed that both forced *hmga1a* expression and forced *hmga1b* expression could suppress hydroxyurea-driven cell death with high statistical significance.

**Figure 6.**
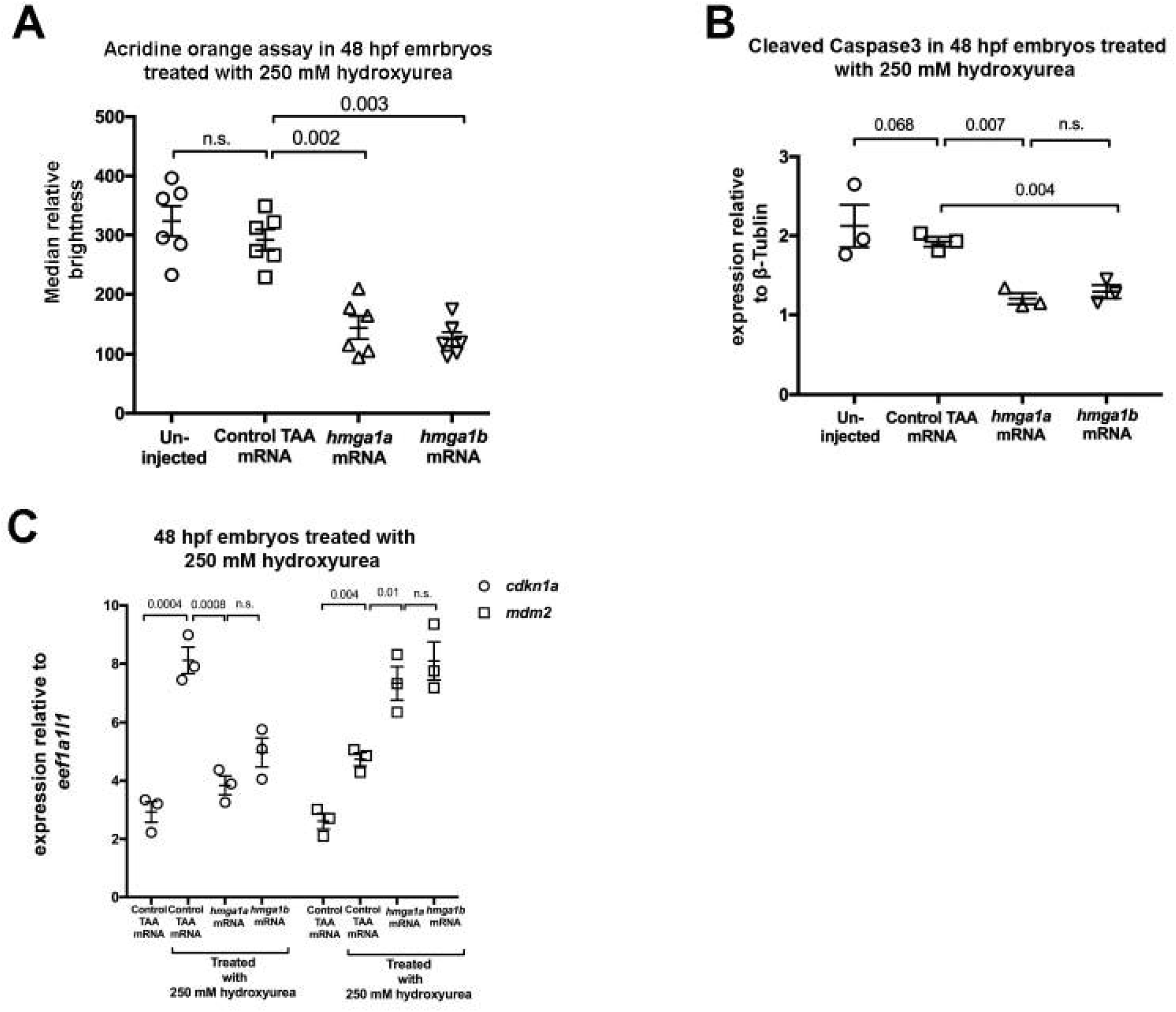
Injection of embryos with *hmga1a* or *hmga1b* mRNA decreases p53-dependent apoptosis. A) Embryos injected with *hmga1a* or *hmga1b* mRNA and treated with 250 mM hydroxyurea show decreased acridine orange staining indicating decreased cell death. B) Embryos injected with *hmga1a* or *hmga1b* mRNA and treated with 250 mM hydroxyurea show decreased cleaved Caspase-3 levels by western immunoblotting in comparison to injection with non-translatable Control TAA mRNA. C) Embryos injected with *hmga1a* and *hmga1b* mRNA and treated with 250 mM hydroxyurea show altered expression of genes related to p53-dependent apoptosis (*cdkn1a* and *mdm2*) when assayed by qPCR. *P*-values for two-tailed T-tests (unequal variances) for bracketed comparisons are shown. n.s., not statistically significant.

In the retracted paper by Pierantoni et al.^7^ it was reported that increasing dosage of a transgene expressing human isoform HMGA1b could increase p53-stimulated transcription of the gene *MDM2* and suppress p53-stimulated transcription of gene *CDKN1A* (previously known as *WAF1* or *p21*) in a transfected cell culture system. Consistent with this, we found that forced expression of either Hmga1a or Hmga1b by mRNA injection into zebrafish zygotes was able to suppress hydroxyurea-driven induction of transcription of the p53-activated gene *cdkn1a* and to increase the induction by hydroxyurea of transcription of the gene *mdm2* (Figure 6C). These data support the idea of an intimate interaction between Hmga1a/b and p53 in the regulation of cell death in vertebrates.

## DISCUSSION

The human *HMGA1* gene has been studied substantially in relation to tumour progression and other disease. To gain better insight into *HMGA1* function and dysfunction, various animal models have been used. The zebrafish and its embryos are a highly manipulable vertebrate model for genetic studies due to a combination of advantageous features including easy embryo accessibility, large zygote size, relatively constant embryo size during its rapid development, short generation time and a well-characterised genome ^32^. We have previously identified and studied a zebrafish co-orthologue of this gene, *hmga1a ^3^*.

The zebrafish *hmga1a* and *hmga1b* genes are co-orthologues of human *HMGA1*. They are duplicate genes that arose from an ancestral gene through the whole-genome duplication early in the evolution of the teleost lineage ^22^. The structure of the putative protein coded by *hmga1b* differs significantly from that of *hmga1a* in the third (downstream) nucleic acid-binding AT-hook domain. This has diverged significantly from the consensus sequence preserved in both human HMGA1a and HMGA1b isoforms and zebrafish Hmga1a. In that sense, and in the lack of the C-terminal acidic tail domain, the putative protein product of *hmga1b* resembles somewhat the human HMGA1c isoform. This suggests subfunctionalisation of *hmga1b* to maintain functions similar to those of HMGA1c and specific to this isoform. However, we note that the C-terminal of the Hmga1b protein consists of two acidic glutamine residues so we cannot yet state with confidence that its function in transcription control differs greatly from the acidic tail domain of Hmga1a.

The lack of conservation of the most C-terminal AT-hook domain in Hmga1b protein affects it ability to control alternative splicing of *psen1* transcripts to produce the PS1IV isoform that is equivalent to the PS2V isoform produced from human PSEN2. Unlike Hmga1a, the Hmga1b protein does not appear able to promote PS1IV formation. This implies that the HMGA1c isoform may not affect alternative splicing of *PSEN2* transcripts to generate PS2V in human cells, although this has not been tested experimentally. However, the structural difference between the zebrafish Hmga1a and Hmga1b co-orthologous proteins does not appear to affect their influence on the p53-regulated genes *cdkn1* and *mdm2*. This is also consistent with the data in the retracted Pierantoni et al paper ^7^ where a truncated form of HMGA1b lacking the downstream, third AT-hook domain retained the ability to bind to p53.

*HMGA1* orthologous genes are expressed during vertebrate embryogenesis ^3,33–35^ and are also commonly upregulated during cancer progression ^28^. They are upregulated in response to hypoxia via a mechanism dependent on reactive oxygen species ^9,27^. We observed a peak of *hmga1b* transcript levels at 48 hpf, the time at which zebrafish embryogenesis ends with hatching from the chorion. This is consistent with a possible restriction of oxygen flow caused by the presence of the chorion during fish embryo development ^36^.

The differences and similarities in structure and function between the two zebrafish co-orthologues of human *HMGA1* are an excellent illustration of how such ancient duplicate genes can be serendipitous tools for dissection of gene/protein activity. This, combined with the ease of exposure of zebrafish embryos to hypoxia and drugs and their facility as an oncogenesis model ^37^ justify more extensive investigation of HMGA1 functions in zebrafish.

## Acknowledgements

This work was supported by the Australian National Health and Medical Research Council (NHMRC, GNT 1061006 for M.L. and M.N., and GNT 1045507 for S.H.M.N). Access to Genomatix licensed software was enabled by a Flinders University 2017 Equipment Grant from the Faculty of Medicine, Nursing and Health Sciences (A.L.).

## Authors Contributions

Designed the experiments and supervised the study (S.H.M.N and M.L). Performed the experiments (S.H.M.N., M.N., A.L., and T.J). Wrote the paper (M.L., and S.H.M.N). Helped review the manuscript (M.L., M.N, and A.L).

## Competing Interests

The authors declare that they have no competing interests.

